# Insights into substrate coordination and glycosyl transfer of poplar cellulose synthase-8

**DOI:** 10.1101/2023.02.07.527505

**Authors:** Preeti Verma, Albert L. Kwansa, Ruoya Ho, Yaroslava G. Yingling, Jochen Zimmer

## Abstract

Cellulose is an abundant cell wall component of land plants. It is synthesized from UDP-activated glucose molecules by cellulose synthase, a membrane-integrated processive glycosyltransferase. Cellulose synthase couples the elongation of the cellulose polymer with its translocation across the plasma membrane. Here, we present substrate and product-bound cryogenic electron microscopy structures of the homotrimeric cellulose synthase isoform-8 (CesA8) from hybrid aspen (poplar). UDP-glucose binds to a conserved catalytic pocket adjacent to the entrance to a transmembrane channel. The substrate’s glucosyl unit is coordinated by conserved residues of the glycosyltransferase domain and amphipathic interface helices. Site-directed mutagenesis of a conserved gating loop capping the active site reveals its critical function for catalytic activity. Molecular dynamics simulations reveal prolonged interactions of the gating loop with the substrate molecule, particularly across its central conserved region. These transient interactions likely facilitate the proper positioning of the substrate molecule for glycosyl transfer and cellulose translocation.

**Highlights:**

- Cryo-EM structures of substrate and product bound poplar cellulose synthase provide insights into substrate selectivity
- Site directed mutagenesis signifies a critical function of the gating loop for catalysis
- Molecular dynamics simulations support persistent gating loop – substrate interactions
- Gating loop helps in positioning the substrate molecule to facilitate cellulose elongation
- Conserved cellulose synthesis substrate binding mechanism across the kingdoms

## Introduction

Cellulose is an abundant biopolymer that is produced primarily by land plants as a structural cell wall component. Because plants produce cellulose from photosynthetically synthesized glucose molecules, the polysaccharide is a major atmospheric carbon dioxide sink as well as a significant renewable energy resource (Carroll and Somerville, 2009).

Cellulose’s glucosyl units are connected via β-(1,4)-glycosidic linkages that enable an approximately 180-degree rotation of neighboring sugar units within the polymer (Nishiyama et al., 2003). The resulting amphipathic polysaccharide can be organized into cable-like fibrillar structures, so-called cellulose micro- and macrofibrils, that are spun around the cell as a load-bearing wall component (Yang and Kubicki, 2020). Cellulose is synthesized from UDP-activated glucose (UDP-Glc) by cellulose synthase (CesA), a membrane-integrated processive family-2 glycosyltransferase (GT) (McNamara et al., 2015; Turner and Kumar, 2018).

CesA catalyzes glucosyl transfer from UDP-Glc (the donor sugar) to the C4 hydroxyl group at the non-reducing end of the nascent cellulose polymer (the acceptor). Following chain elongation, CesA also facilitates cellulose translocation across the plasma membrane through a pore formed by its own transmembrane (TM) segment. To couple cellulose synthesis with secretion, CesA’s catalytic GT domain packs against a channel-forming TM region via three conserved amphipathic interface helices (IF1-3) (Morgan et al., 2013; Morgan et al., 2016).

Cellulose biosynthesis is evolutionarily conserved, with homologous pathways found in prokaryotes, oomycetes, and some animals. Previous work on bacterial cellulose biosynthetic systems from *Gluconacetobacter xylinum* (formerly *Acetobacter xylinus*) (Brown et al., 1976; Du et al., 2016), *Rhodobacter sphaeroides (Omadjela et al*., *2013)*, and *Escherichia coli* (Bokranz et al., 2005) provided detailed insights into the reaction mechanism and enzyme regulation (Fang et al., 2014; Morgan et al., 2014; Richter et al., 2020; Ross et al., 1987), as well as cellulose secretion (Morgan *et al*., 2016), assembly (Abidi et al., 2022; Nicolas et al., 2021), and modification (Thongsomboon et al., 2018). Further, recent cryogenic electron microscopy (cryo-EM) studies on trimeric plant CesA complexes confirmed an evolutionarily conserved enzyme architecture, in support of an equally conserved catalytic reaction mechanism (Purushotham et al., 2020; Zhang et al., 2021).

To delineate principles underlying substrate selectivity and catalysis, we determined cryo-EM structures of the full-length poplar CesA isoform-8 (CesA8) bound to either UDP-Glc, or the product and competitive inhibitor UDP. The obtained complexes demonstrate substrate binding to a conserved pocket at the interface between the enzyme’s cytosolic catalytic domain and TM region. However, in contrast to the bacterial homolog BcsA, a conserved ‘gating loop’ that stabilizes UDP-Glc at the active site (Morgan *et al*., 2016), is disordered in the substrate-bound CesA8 complex. Extensive mutagenesis analyses of the loop’s conserved FxVTxK motif in *Rhodobacter sphaeroides* BcsA as well as poplar CesA8 underscore its importance for catalytic activity. All-atom molecular dynamics simulations indeed confirm prolonged interactions of the loop with the substrate molecule and suggest its role in proper positioning of the substrate for glycosyl transfer.

## Results

### Cryo-EM analyses of nucleotide-bound poplar CesA8

Poplar CesA8 was expressed in Sf9 insect cells and purified in the detergents lauryl maltose neopentyl glycol (LMNG)/cholesteryl hemisuccinate (CHS) and glyco-diosgenin (GDN) as previously described (Purushotham *et al*., 2020) and summarized in the Star Methods. Under these conditions, the enzyme is catalytically active, synthesizing cellulose *in vitro* in the presence of UDP-Glc and magnesium ions. Alongside cellulose, CesA generates UDP as a second reaction product of the glycosyl transfer reaction. Previous studies on bacterial and plant cellulose synthases as well as other related family-2 GTs demonstrated that UDP competitively inhibits the enzymes, due to its interactions with the catalytic pocket (Kumari and Weigel, 1997; Omadjela *et al*., 2013; Purushotham et al., 2016). This observation has been exploited to obtain UDP-inhibited structures of hyaluronan and cellulose synthases (Maloney et al., 2022; Morgan *et al*., 2014).

We determined UDP and UDP-Glc bound poplar CesA8 structures by incubating the purified enzyme with 5 mM of nucleotide and 20 mM MgCl_2_ prior to cryo grid preparation (see Star Methods, Figure S1). The ligand bound CesA complexes were imaged and processed as described before (Purushotham *et al*., 2020) (Figure S1-2 and Table S1). Overall, the trimeric organization of CesA8 is preserved in UDP and UDP-Glc bound states, suggesting that the complex indeed represents a biologically functional unit (Figure 1A). Within the resolution limits of our cryo-EM maps (approximately 3.5 Å), the UDP moiety adopts the same binding pose in the UDP-only and UDP-Glc bound states (Figure S3A), hence, the following discussion focuses on the substrate-bound conformation. Each CesA8 protomer also contains a nascent cellulose polymer within the TM channel. The polymer’s first five glucosyl units, starting at the non-reducing end near the catalytic pocket, are sufficiently well ordered to allow modeling (Figure S2). As described previously (Purushotham *et al*., 2020), the terminal acceptor glucosyl unit rests next to Trp718 of the conserved QxxRW motif, right above the substrate binding pocket (Figure 1B).

**Figure 1.**
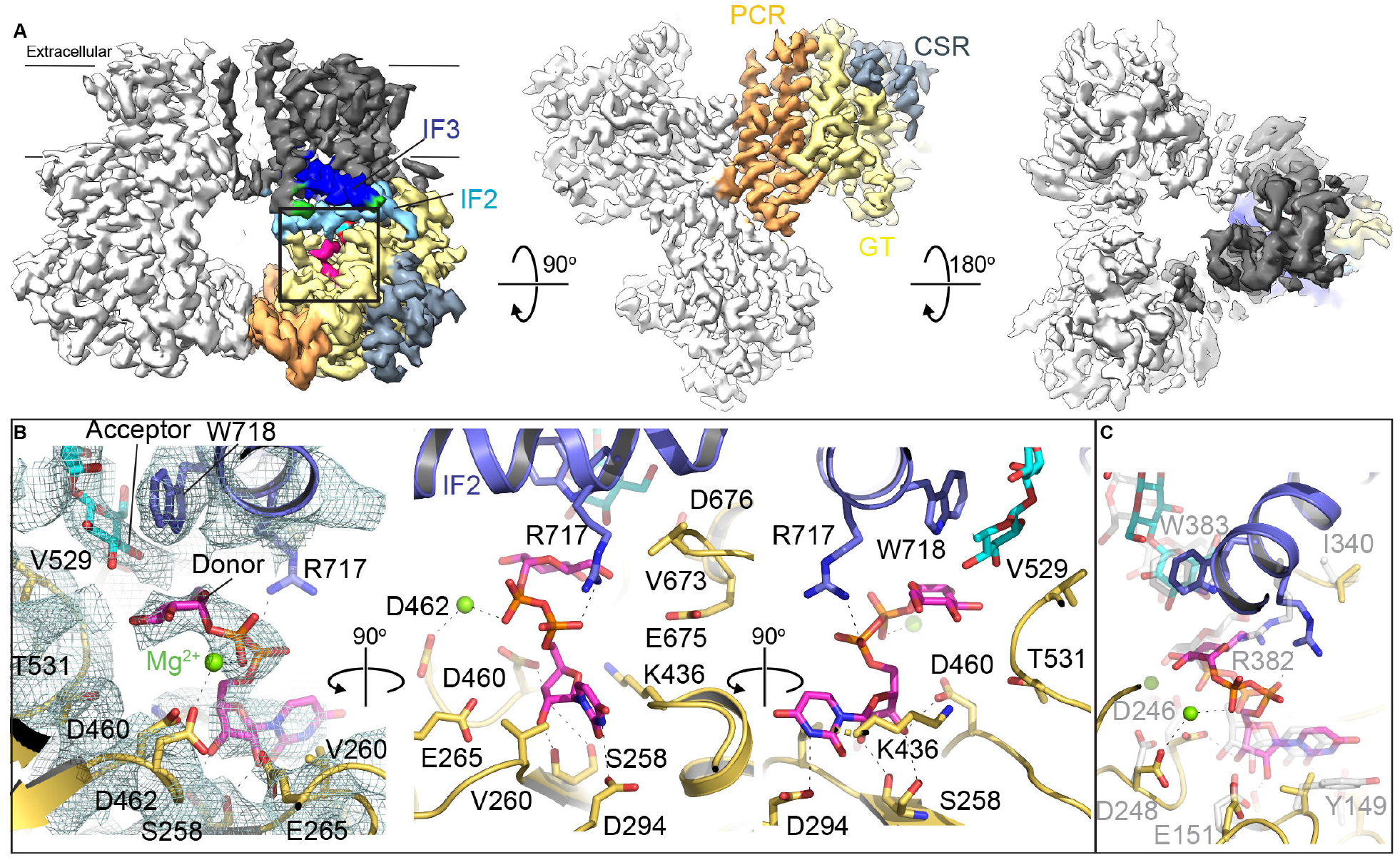
Structure of UDP-glucose bound poplar CesA8. **(A)** Cryo-EM map of the substrate-bound poplar CesA8 trimer. Two subunits are shown in light gray and one subunit is colored according to its domains: dark gray: TM region, yellow: GT domain, wheat: PCR, steelblue: CSR, and lightblue and blue for IF helices 2 and 3, respectively. See also Figure S1-S2, S3A. **(B)** Zoom- in view of the active site and substrate coordination. The cryo-EM map of the substrate and surrounding residues is shown as a mesh in the left panel. **(C)** Superimposition of substrate-bound BcsA (PDB: 5EIY) and poplar CesA8. BcsA is colored light gray for its carbon atoms. See also Figure S3.

### CesA8 positions the donor sugar beneath the acceptor glucosyl unit

CesA8’s GT domain forms a classical GT-A fold with a central mixed β-sheet surrounded by α-helices (Lairson et al., 2008). The bound substrate molecule is coordinated by conserved residues distributed throughout the GT domain (Morgan *et al*., 2013) (Figure 1B). First, the substrate’s uridine group is sandwiched between Glu265 and Lys436 and fits into a groove created by Ser258 and Val260 of the conserved STVDP motif belonging to the first β-strand of the GT-A fold. Second, Asp294 of the invariant DDG motif terminating β-strand #2 is in hydrogen bond distance to the Nε ring nitrogen of the uracil moiety. Third, the conserved DxD motif (Asp460 and Asp462), following β-strand #5, contributes to the coordination of a magnesium cation, which is also in contact with the substrate’s β-phosphate. Additionally, the nucleotide’s diphosphate group interacts with Arg717 of the QxxRW motif originating from IF-2 (Figure 1B).

The donor sugar fits into a polar pocket directly underneath the acceptor glucosyl unit of the nascent cellulose chain. This pocket is proximal to the water-membrane interface formed from IF-2, the finger helix that is N-terminally capped with the invariant VTED motif (residues 673 to 676), as well as the backbone of Val529, Gly530 and Thr531 belonging to the conserved YVGTG motif (Figure 1B). Potential hydrogen bond donors and acceptors from protein side chain and backbone regions surround the donor glucosyl unit. However, all observed distances to the donor’s hydroxyl groups exceed 3.5 Å, suggesting that the substrate molecule is not fully inserted into the catalytic pocket. Accordingly, the distance between the acceptor’s C4 hydroxyl and the donor’s C1 carbon exceeds 5.5 Å (Figure 1B). The observed substrate coordination is consistent with interactions delineated for bacterial BcsA bound to a non-hydrolysable UDP-Glc phosphonate analog (Morgan *et al*., 2016) (Figure 1C), as well as UDP-N-acetylglucosamine-bound to chitin and hyaluronan synthases (Chen et al., 2022; Maloney *et al*., 2022; Ren et al., 2022) (Figure S3B-D).

### The flexible gating loop is required for catalytic activity

CesA8’s IF3 is connected to TM helix 5 via a ∼20 residue long cytosolic gating loop that runs roughly across the opening of the catalytic pocket. The gating loop contains a conserved FxVTxK motif but is not resolved in all cryo-EM maps of CesAs, most likely due to conformational flexibility (Figure 2A).

**Figure 2.**
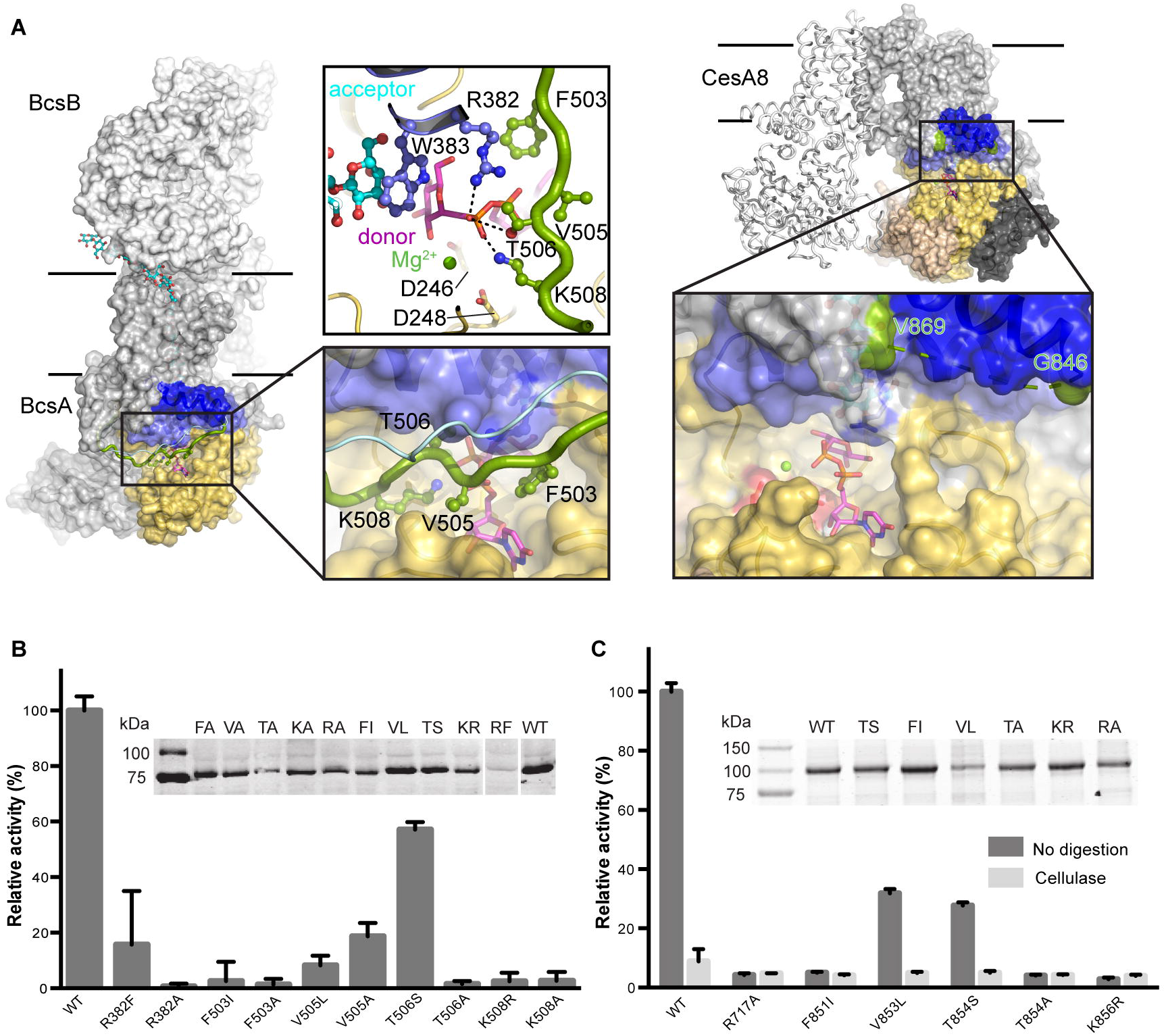
The gating loop is critical for catalytic activity. **(A)** Substrate bound structures of bacterial (BcsA, PDB: 5EIY) and poplar cellulose synthase (this study). The substrate is shown as sticks colored magenta for carbon atoms. The gating loop positions observed in BcsA in the absence (BcsA, PDB: 4P02) and the presence of substrate are shown as light blue and green ribbons, respectively. **(B)** Comparison of the catalytic activities of the bacterial BcsA-B wild-type (WT) complex and its variants containing the indicated BcsA mutants. Experiments were performed in IMVs and activities were measured upon incorporation of tritium-labelled glucose into cellulose. The product yield for the WT enzyme was set to 100%. Inset: Western blot of the BcsA-B containing IMVs used for activity assays detecting His-tagged BcsA. Blotting was done to normalize the concentration of mutants with respect to the WT enzyme. **(C)** Catalytic activity determined for WT CesA8 and its mutants. Reactions were performed in the presence or absence of cellulase using affinity purified proteins. The product obtained for the undigested WT CesA8 was set as 100%. Inset: Coomassie stained 10% SDS-PAGE of the purified CesA8 constructs used for normalization and activity assays. Error bars represents positive standard deviations from means of three replicas. See also Figure S4.

Crystallographic analyses of *Rhodobacter sphaeroides* BcsA revealed different conformations of the gating loop (Morgan *et al*., 2013; Morgan *et al*., 2014). The loop retracts from the catalytic pocket in a nucleotide-free state and inserts into it in the presence of either UDP or a substrate analog. In the inserted state, the conserved FxVTxK motif contacts the UDP moiety, thereby likely stabilizing it at the active site. In addition to substrate stabilization, gating loop insertion into the catalytic pocket also facilitates translocation of the nascent polysaccharide between elongation steps (Morgan *et al*., 2016). Unlike for BcsA, nucleotide binding to CesA8 does not stabilize the gating loop in a similar manner (Figure 2A).

To probe the functional significance of the gating loop’s FxVTxK motif, we performed site-directed mutagenesis and *in vitro* functional analyses of the bacterial BcsA and poplar CesA8 enzymes. For BcsA, replacing Phe503 of the FxVTxK motif with Ala or the bulky hydrophobic residue Ile abolishes catalytic activity (Figure 2B and Figure S4). Substituting the conserved Val505 residue with Ala or Leu dramatically reduces catalytic activity to about 20 and 10%, respectively, relative to the wild-type enzyme. A drastic reduction is observed when the following Thr506 residue is replaced with Ala, yet its substitution with Ser retains about 60% relative catalytic activity. The Lys508 residue at the C-terminus of the FxVTxK motif is also critical for function, neither an Ala nor an Arg residue at this position supports enzymatic activity (Figure 2B).

Upon insertion into the catalytic pocket, Phe503 of BcsA’s gating loop forms cation-π interactions with Arg382 of the conserved QxxRW motif located in IF2 at the cytosolic water-lipid interface (Morgan *et al*., 2014). In this position, Arg382 forms a salt bridge with the substrate’s diphosphate group, as also observed in CesA8 (Figure 1B and 2A). This residue is critical for catalytic activity as its substitution with Ala renders BcsA inactive, while its substitution with Phe retains about 16% activity (Figure 2B).

A similar analysis was performed of CesA8’s FxVTxK motif. Here, we replaced Phe851 with Ile, V853 with Leu, Thr854 with Ser or Ala, and Lys856 with Arg. Due to increased background readings in membrane vesicles, the mutant enzymes were purified and analyzed for catalytic activity in a micelle-solubilized state, as previously described (Purushotham *et al*., 2020). Of the generated mutants, only the V853L and T854S substitutions retain about 30% catalytic activity, relative to the wild-type enzyme. Further, as observed for BcsA, replacing Arg717 of CesA8’s QxxRW motif with Ala renders the enzyme catalytically inactive (Figure 2C).

### Molecular dynamics simulations reveal persistent gating loop – substrate interactions

To probe the interactions between CesA8’s gating loop and a nucleotide at the active site, we examined the dynamics of the gating loop in the presence of UDP and UDP-Glc. The initial conformation of the loop was modeled based on the loop’s position in the substrate-bound BcsA crystal structure (PDB: 5EIY) as well as weak discontinuous gating loop density observed at low contour levels in the CesA8 cryo-EM map (Figure 3 and Figure S5). All-atom molecular dynamics (MD) simulations of UDP- and UDP-Glc-bound CesA8 monomers with a phospholipid bilayer, water, and NaCl were performed for 1,000 ns, as detailed in the Star Methods. Relative to its starting conformation, the gating loop remains in close contact with the nucleotide during the simulations, with greatest fluctuations observed for regions N- and C-terminal to the conserved FxVTxK motif (Figure 3A). For the UDP-bound case, notable contacts are observed for all residues of the FxVTxK motif, except for Thr852 and Lys856. For the UDP-Glc-bound case, a similar set of residues of the FxVTxK motif are found to have influential interactions. In both cases, residues 853 and 854 (i.e., the conserved VT) are predicted to interact the most with UDP or UDP-Glc. Phe851 and Val853 are positioned to surround the substrate’s uracil group, while Thr854 hydrogen bonds with the ligand’s alpha phosphate (Figure 3B) with hydrogen bond times of ∼100% of the total simulated time with both UDP and UDP-Glc (Tables S2-3). The C-terminal Lys856 of the FxVTxK motif is predicted to interact minimally and transiently with the phosphates and the uracil group. The mean contact score metric employed in PyContact provides a measure of contact persistence over time and contact proximity; this metric is based on a distance-weighted sigmoidal function, and the contact scores are averaged over time providing a mean quantity. Mean contact scores associated with the gating loop residues and UDP/UDP-Glc/Mg^2+^ reveal relatively high contact persistence and proximity, especially for the gating loop’s central Val853 and Thr854 (Figure 3C and Tables S2-3).

**Figure 3.**
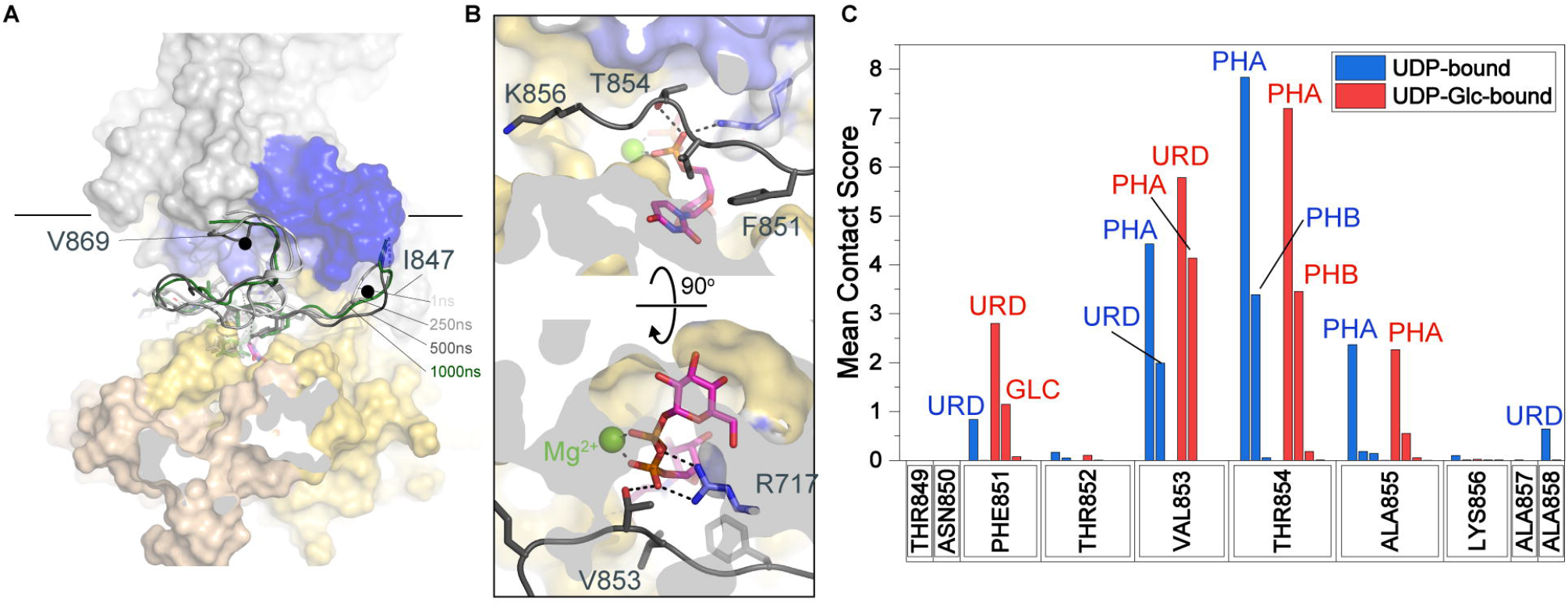
Molecular dynamics simulation analysis of gating loop – UDP interactions. **(A)** Overlay of UDP-Glc bound CesA8 models with an inserted gating loop after 1, 355, 500 and 1000 ns of MD simulation. The original model is shown as a surface-colored structure as in Figure 1. The gating loop is shown as a cartoon for models obtained after the indicated simulation times. **(B)** Close up view of interactions between the gating loop and the substrate’s uracil moiety after 1000 ns of MD simulation. **(C)** A bar plot of mean contact scores for individual CesA gating loop residues interacting with UDP/UDP-Glc and/or Mg^2+^. Generated with Origin (Origin). URD: Uridine, PHA/B: alpha and beta phosphate, MG: magnesium. See also Figure S5.

## Discussion

CesA catalyzes multiple reactions: First, the formation of a linear β-(1,4)-glucan; second, the secretion of the polysaccharide into the extracellular milieu; and third, due to the self-assembly of CesAs into supramolecular complexes, the coalescence of cellulose polymers into fibrillar structures (McNamara *et al*., 2015; Turner and Kumar, 2018).

Polymer secretion is achieved by closely associating the catalytic cytosolic GT domain with a channel-forming TM segment, with residues from both regions contributing to donor and acceptor coordination.

The observed binding pose of the substrate UDP-Glc at CesA8’s active site is consistent with other structures of processive and non-processive GT-A enzymes, including hyaluronan and chitin synthases (Chen *et al*., 2022; Maloney *et al*., 2022; Ren *et al*., 2022) (Figure 4A and B). CesA coordinates UDP with invariant sequence motifs, primarily localized at the edge of its central GT-A β-sheet. The donor sugar, attached to UDP, is positioned in a hydrophilic pocket near the water-lipid interface. This pocket is created by the GT domain as well as amphipathic interface helices that establish the transition from the cytosolic to the TM region. The observed substrate coordination contrasts a recently reported UDP-Glc-bound crystal structure of the GT core of *Arabidopsis* CesA3 (Qiao et al., 2021), which exhibits an unphysiological catalytic pocket.

**Figure 4.**
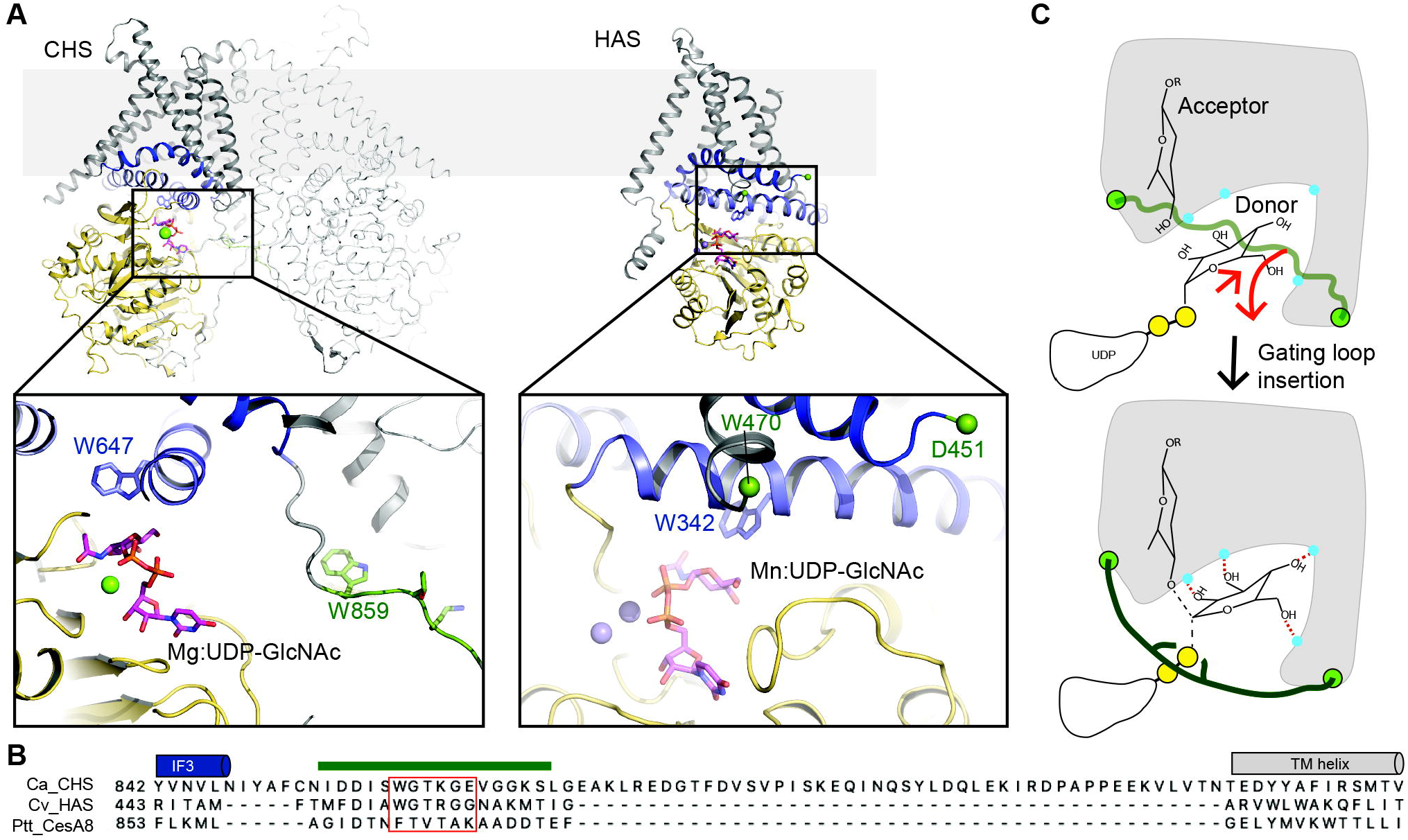
Localization and proposed function of the gating loop in membrane integrated processive GT-2 glycosyltransferases. **(A)** Substrate bound structures of *Candida albicans* chitin synthase (CHS, PDB: 7STM) and Chlorella virus hyaluronan synthase (HAS, PDB: 7SP8). Both enzymes place a conserved WGTR/KG motif near the catalytic pocket (unresolved in HAS). **(B)** Alignment of the gating loop regions of chitin, hyaluronan and cellulose synthases. **(C)** Gating loop insertion likely positions the substrate molecule closer to the base catalyst and helps in accepting glycosyl unit. Conformational changes of the donor sugar may increase the reactivity of the electrophile.

Compared to UDP or substrate-bound states of bacterial BcsA, CesA8’s conserved gating loop remains disordered in the substrate-bound cryo-EM structure. Weak map density ‘above’ the entrance to the catalytic pocket (Figure S5A) indicates flexibility of the gating loop, as also observed for BcsA in nucleotide-free states (Morgan *et al*., 2014). Site-directed mutagenesis of the gating loop, however, demonstrates its profound importance for cellulose biosynthesis. We hypothesize that the gating loop inserts transiently into the catalytic pocket to position the substrate for glycosyl transfer and perhaps decreasing the distance between the donor and acceptor glucosyl units (Figure 4C). Substrate repositioning may also (a) favor protein interactions with the donor’s hydroxyl groups to foster substrate specificity, and (b) facilitate conformational changes of the donor’s pyranose ring to increase the reactivity of its C1 carbon.

Hyaluronan and chitin synthases also contain putative gating loops. The corresponding sequence is WGTR/KG instead of FxVTxK (Figure 4B). This motif is located in a loop following IF3 near the active site. It is disordered in hyaluronan synthase structures or bridges a dimerization motif in chitin synthase (Figure 4A). Substrate-bound structures of these enzymes also suggest incomplete insertions of the substrate molecules into the catalytic pockets, based on the assumed acceptor binding site formed by the Trp of the conserved QxxRW motif. As for CesA, transient gating loop insertion could enforce a substrate conformation favorable for glycosyl transfer (Figure 4C).

Cellulose microfibrils are synthesized from CesA complexes (CSC) resembling six-fold symmetric particles (Nixon et al., 2016). The repeat unit is likely represented by the trimeric cryo-EM structures obtained for poplar and cotton CesAs (Purushotham *et al*., 2020; Zhang *et al*., 2021). Within a CSC, multiple CesAs synthesize and secrete cellulose polymers to facilitate their alignment into a microfibril. It is currently unknown whether the individual CesA activities are coordinated. Our substrate-bound cryo-EM structures do not suggest allosteric regulation of the catalytic activities within a CesA trimer. However, we cannot exclude interprotomer crosstalk in the context of a fully assembled CSC, perhaps mediated by currently unresolved regions.

## Acknowledgements

The authors thank Balasubramanyam Chittoor for initial cloning of CesA8 mutants and purifications. We are grateful to the staff at the EM core facilities at the University of Virginia (MEMC, Kelly Dryden and Michael Purdy) as well as the Laboratory for Biomolecular Structure (LBMS) at Brookhaven National Laboratory (Liguo Wang). LBMS is supported by the DOE Office of Biological and Environmental Research (KP1607011). MD simulations by A.L.K. and Y.G.Y. and work on poplar CesA8 by R.H. and J.Z. were supported by the Center for Lignocellulose Structure and Formation, an Energy Frontier Research Center funded by the U.S. Department of Energy, Office of Science, Basic Energy Sciences (award DESC0001090). Work on bacterial BcsA performed by P.V. and J.Z was supported by NIH grant R35GM144130 awarded to J.Z. J.Z. is an investigator of the Howard Hughes Medical Institute. This article is subject to HHMI’s Open Access to Publications policy. HHMI lab heads have previously granted a nonexclusive CC BY 4.0 license to the public and a sublicensable license to HHMI in their research articles. Pursuant to those licenses, the author-accepted manuscript of this article can be made freely available under a CC BY 4.0 license immediately upon publication.

## Author Contributions

P.V. performed all mutagenesis experiments, R.H. prepared cryo grid samples and collected all EM data. J.Z. processed the data and built the models. A.L.K. and Y.G.Y. performed the MD analysis. All authors evaluated the data. J.Z. wrote the initial manuscript and all authors edited the manuscript.

## STAR⍰Methods

### Resource availability

#### Lead contact

Further information and request for resources and reagents should be directed to and will be fulfilled by the lead contact, Jochen Zimmer.

#### Materials availability

This study did not generate new unique reagents. Plasmids generated in this study will be available upon request to the lead contact.

#### Data and code availability

- The coordinates of UDP bound CesA8 and UDP-Glucose bound CesA8 have been deposited to the Protein Data Bank under accession numbers XXX and YYY. CryoEM maps of the UDP bound CesA8 and UDP-Glucose bound CesA8 have been deposited in the Electron Microscopy Data Bank under the accession numbers XXXX and YYYY respectively. All the deposited data are publicly available as of the date of publication and accession numbers are also listed in the key resources table.
- This paper analyzes existing, publicly available data. These accession numbers for the datasets are listed in the key resources table.
- This paper does not report original code data.
- Any additional information required to reanalyze the data reported in this paper is available from the lead contact upon request.

### Experimental model and subject details

#### Bacterial strains

Escherichia coli Rosetta 2 (DE3) cells (Novagen) were used in this study for recombinant protein production. Cells were cultured in ZYP-5052 auto-induction media (Studier, 2005) supplemented with necessary antibiotics.

#### Cell lines

Hybrid Aspen (poplar) CesA8 was expressed in *Spodoptera frugiperda* (Sf9) cells infected with recombinant baculovirus (pACEBac1). SF9 cells were grown in ESF921 medium (Expression systems) at 27°C and cells were harvested 72 h after the infection when cell viability was dropped down to 70%.

### Method details

#### Mutagenesis

The BcsA gating loop (F503A/I, V505A/L, T506A/S, K508A/R) and R382A/F mutants were generated from the wild-type (WT) construct as described earlier (Morgan *et al*., 2016) via QuikChange mutagenesis. The WT type construct contains both the *Rhodobacter sphaeroides* cellulose synthase (bcs)A and bcsB genes expressed in pETDuet-1 vector, wherein BcsA was expressed with a C-terminal dodeca-histidine tag. Similarly, the hybrid aspen CesA8 mutants were generated from an existing WT pACEBac-CesA8 plasmid (Purushotham *et al*., 2020) by the same method. Not all mutations generated for BcsA were also introduced into CesA8. The CesA8 mutants were designed based on activity results obtained for the BcsA mutants.

All the oligonucleotides used in generating the BcsA and CesA8 mutants are provided in Table S5.

#### Expression and purification of CesA8 and its variants in SF9 insect cells

Expression of CesA8 and its mutants was performed in SF9 insect cells and the purification was carried out as described previously (Purushotham *et al*., 2020), with some modifications. Briefly, the cell pellet from 1-1.5 L of culture was resuspended in the modified buffer [Buffer A: 20 mM Tris pH 7.5, 100 mM NaCl, 5 mM sodium phosphate, 5 mM sodium citrate, 1 mM TCEP] containing the detergents 1% lauryl maltose neopentyl glycol (LMNG, Anatrace) and 0.2% cholesteryl hemisuccinate (CHS, Anatrace), supplemented with protease inhibitor cocktail (0.8 μM Aprotinin, 5 μM E-64, 10 μM Leupeptin, 15 μM Bestatin-HCl, 100 μM AEBSF-HCl, 2 mM Benzamidine-HCl and 2.9 mM Pepstatin A). The entire mixture was lysed with 20 strokes of a tight fitting 100 mL dounce homogenizer. After solubilization at 4°C for 1 hour, the insoluble material was removed by centrifugation at 42,000 rpm for 45 min in a Ti45 rotor. All subsequent steps including the exchange of detergent from LMNG-CHS to 0.02% glycol-diosgenin (GDN, Anatrace) during the wash buffers were carried out as described before (Purushotham *et al*., 2020), except for the omission of size exclusion chromatography. Instead, a dialysis step was performed wherein the Ni-NTA eluent was concentrated to approximately 5 mL using a 100-kDa spin concentrator (Millipore) and then dialyzed overnight against Buffer A containing 0.02% GDN. The following day, the dialyzed protein was concentrated to roughly 1.6-1.9 mg/mL and flash-frozen in small aliquots in liquid nitrogen and then stored at -80°C until further use.

#### Expression and Preparation of Inverted Membrane Vesicles of Rhodobacter sphaeroides BcsA-B complex and its variants

Expression and inverted membrane vesicle (IMV) preparation was carried out as described previously (Morgan *et al*., 2013; Omadjela *et al*., 2013) for the wild-type BcsA-B complex. Briefly, the BcsA-B complex was expressed in *Escherichia coli* Rosetta 2 cells in auto-induction medium. The 2 L cell pellet obtained for each protein construct was resuspended in Resuspension Buffer (RB) containing 20 mM Tris pH 7.2, 100 mM NaCl, and 10% Glycerol, supplemented with 1 mM phenylmethylsulfonyl fluoride (PMSF). Cells were lysed in a microfluidizer followed by centrifugation at 12,500 rpm in a Beckman JA-20 rotor for 20 min. The supernatant was carefully recovered and roughly 25 ml was layered over a 1.8 M sucrose cushion made in RB buffer, followed by centrifugation at 42,000 rpm for 120 min in a Ti45 rotor. The dark brown ring formed at the sucrose cushion was carefully withdrawn, diluted five-fold in the RB buffer, and finally, the membrane vesicles were sedimented via centrifugation at 42,000 rpm for 90 min in a Ti45 rotor. The pellet fraction was rinsed with RB buffer, resuspended in 1 ml RB, and homogenized using a no. 6 paintbrush followed by douncing in a 2 ml grinder. The vesicles were aliquoted in small quantities and flash frozen in liquid N_2_ until further use. All the steps were performed at 4ºC.

#### Normalization of protein levels for activity assays

To normalize the concentration of BcsA amongst the WT and mutant IMVs for performing the activity assays, freeze-thawed IMVs were treated with 2% SDS to aid in solubilization and proper migration during SDS-PAGE. Gel loading samples were made from these solubilized IMVs and western blotting against the BcsA-His tag was performed. The band intensities after Western blotting were used to calibrate the amounts of IMVs used for activity assays.

CesA8 protein concentrations for WT and its mutants were normalized based on quantitative SDS-PAGE analysis of the purified proteins. The gel was imaged after Coomassie-staining and further analyzed using LI-COR Odyssey Imaging system to compare the band intensities.

#### In vitro cellulose synthesis

For *Rhodobacter* BcsA-B, WT and mutant IMVs were used for assaying cellulose biosynthetic activity as described (Omadjela *et al*., 2013). In the beginning, a time course was conducted using the wild-type BcsA-B IMVs to find an incubation time in the linear phase of product accumulation. Synthesis reaction was performed by incubating IMVs with 5 mM UDP-Glucose and 0.25 µCi of UDP-[^3^ H]-glucose, 30 µM cyclic-di-GMP (c-di-GMP) and 20 mM MgCl_2_ in RB buffer lacking glycerol. Here, cellulose biosynthesis reaction was monitored for different time periods starting from 0 to 180 minutes. Aliquots were spotted onto Whatman-2MM chromatography paper and developed by descending paper chromatography using 60% ethanol. The polymer retained at the origin was quantified by scintillating counting. Based on this time course, all standard reactions were carried out at 37ºC for 30 min. All reactions were performed in triplicate and error bars represent deviations from the means.

For CesA8, the purified, micelle solubilized protein was used for synthesis reactions. The assays were performed as described previously (Purushotham *et al*., 2020). The activity assay conditions were same as for BcsA-B but with no c-di-GMP and in Buffer A containing 0.02% GDN. As for assays with the bacterial enzyme, we determined a suitable assay time point during the linear product accumulation. Based on this time course, all subsequent reactions were performed for 30 min at 30ºC. To confirm the formation of authentic cellulose, enzymatic degradations of the *in vitro* synthesized glucan were performed using a commercial cellulase (endo-(1,4)-β-glucanase E-CELTR; Megazyme), wherein 5U of the enzyme was added at the beginning of synthesis reaction. All reactions were performed in triplicate and error bars represent deviations from the means.

#### Cryo-EM data collection

Ligand bound CesA8 complexes were generated by adding 20 mM MgCl_2_ and 5 mM UDP or UDP-Glc to the purified protein and incubation for 30 min on ice prior to cryo grid preparation. Cryo-EM analyses were performed as described before (Purushotham *et al*., 2020). In short, 2.5 µL aliquot was applied to a glow-discharged (in the presence of amylamine) C-flat 400 mesh 1.2/1.3 holey carbon grid (Electron Microscopy Sciences), blotted with Vitrobot Mark IV (FEI, Thermo Fisher Scientific) with force 7 for 12-14 s at 4°C, 100% humidity, and flash frozen in liquid ethane. Grids were screened in-house for optimal ice thickness and particle distribution. High quality data sets were collected at the Brookhaven National Laboratory for BioMolecular Structure (LBMS) on a Titan Krios G3i equipped with a X-FEG electron source, Gatan K3 direct electron detector, and BioQuantum energy filter. Movies were collected in super-resolution mode with a pixel size of 0.4125 and 0.88 Å for the UDP-Glc and UDP-bound complexes, respectively. All movies were collected in counting mode at a magnification of 105,000k and 81K, respectively, and defocus range from -2.3 to -0.8 µm, with a total dose of 51e-/Å2.

#### Data processing

Cryo-EM data processing followed a similar workflow in cryoSPARC (Punjani et al., 2017) as previously described (Purushotham *et al*., 2020). Movies were full-frame motion corrected followed by CTF estimation. Exposures were manually curated based on estimated resolution, defocus, and drift as well as ice contamination.

Initial templates for particle picking were generated using ‘blob picking’ with inner and outer particle diameters of 200 and 350 Å. Particles were extracted with a box size of 600 pixel and Fourier cropped to a box size of 150 pixel. Following 2D classification, selected class averages were used for template-based particle picking. The new particle stack was inspected, extracted with 4-fold Fourier cropping, and classified in 2 and 3-dimensions. The best particles of the UDP-Glc bound dataset were re-extracted using a 640 pixels box and Fourier cropped to a 320 pixels box. For the UDP complex, final particle stack was extracted at a box size of 400 pixels without cropping. Refinements followed standard non-uniform refinement with C3 symmetry.

#### Model building

PDB entry 6WLB was used as an initial model. The previous CesA8 trimer structure was rigid body docked into the EM map in Chimera (Pettersen *et al*., 2004) and manually adjusted in Coot (Emsley and Cowtan, 2004). UDP and UDP-Glc were placed and manually refined in Coot. The model was refined in phenix:refine (Adams et al., 2010) without imposing NCS symmetry. Coordinates and EM maps have been deposited at the ProteinDataBank under accessing codes 8G27 and 8G2J.

#### Molecular dynamics system construction

The MD workflow started from the cryo-EM structure of poplar CesA8 with a bound UDP, coordinated Mg^2+^, and cellopentaose (Purushotham et al., 2020). MD simulations were performed with a monomeric CesA8 construct. Rather than a TM7 (932-958) of the same CesA8 monomer, this monomeric construct contained the TM7 of the adjacent CesA8 subunit to complete its TM channel architecture. Then, the unresolved gating loop was modeled by analogy to BcsA (PDB: 5EIY), representing an inserted state of the gating loop. Subsequently, the SWISS-MODEL web server was used to generate initial coordinates for the remaining residues of the gating loop (847-848 and 856-865); SWISS-MODEL employs the ProMod3 homology modeling engine, which uses a database of structural fragments derived from the Protein Data Bank or falls back to a Monte Carlo approach to generate coordinates (Biasini et al., 2013; Studer et al., 2021; Waterhouse et al., 2018). A UDP-Glc-bound case was also prepared for MD simulation, which involved replacing the UDP molecule with UDP-Glc while remaining consistent with the cryo-EM determined UDP-Glc-bound CesA8 structure. The AMBER 2021 software package (Case *et al*., 2021) was used for all subsequent system construction, force field implementation, and simulation tasks. The simulation environment was assembled around each initial model (CesA8 monomer with UDP/UDP-Glc, Mg^2+^, and cellopentaose). Specifically, AMBER’s packmol-memgen, with Packmol 18.169, was used to add a homogeneous dioleoylphosphatidylcholine (DOPC) phospholipid bilayer, water, and 0.15 M NaCl; unless indicated, default settings such as the padding distances for the lipid bilayer and water around the solute were used (Martinez *et al*., 2009) (Schott-Verdugo and Gohlke, 2019).

#### MD force fields

AMBER’s tleap was then used to apply selected force fields (set of potential energy expressions, parameters, and structural libraries) to the constructed system. The following force fields were used: ff14SB (CesA protein) (Maier et al., 2015), Lipid17 (DOPC) (Case *et al*., 2021; Dickson et al., 2014), GLYCAM06j (cellopentaose) (Kirschner et al., 2008), TIP3P model (water) (Jorgensen et al., 1983), and the 12-6-4 Lennard-Jones (LJ) set of the Li-Merz monovalent and divalent ion parameters for TIP3P (Na^+^, Cl^-^, Mg^2+^) (Li and Merz, 2014, Li et al., 2015). Base parameters for the uridine and phosphate groups of UDP were provided through AMBER’s “parm10.dat”, which includes the OL3 parameters for RNA (Zgarbova et al., 2011). Parameters for the glucosyl group of UDP-Glc were provided through GLYCAM06. The parameters describing the connection between the beta phosphate and the glucosyl group were assigned by analogy to GLYCAM06; specifically, one bond parameter, four angle parameters, and seven dihedral parameters were added based on methyl sulfate, ethyl sulfate, methoxy alkanes, and ether alcohols (Table S4). UDP parameters based on GLYCAM93 have been developed previously (Imberty et al., 1999); while this previous work was used as a general reference point, we sought to remain consistent with the newer GLYCAM06 and thus adopted comparable parameters directly from GLYCAM06. Furthermore, a modification was applied to the 12-6 LJ parameters of the hydroxyl hydrogen atom type “HO” as used in AMBER’s “all_modrna08.frcmod” (Aduri et al., 2007) (Table S4); this was deemed necessary to avoid a known potential issue when using hydroxyl hydrogen atom types with Rmin and epsilon parameters of zero.

#### MD partial charges

For both UDP and UDP-Glc, the net charge was considered as -2.0, that is, a single protonation on the beta phosphate of UDP and no protonation of the UDP-Glc phosphates. This selected protonation state of UDP is based on log(K_a_) values that have been reported for adenosine diphosphate, ADP (7.02, 4.19, and 0.9 for single, double, and triple protonation of the phosphates) via ^1^ H NMR chemical shift data and non-linear regression (Wang et al., 1996). The selected protonation state of UDP-Glc is based on pK_a_ values that have been reported for UDP-GlcNAc – an N-acetyl derivative of UDP-Glc (6.6 and 6.3 for the alpha and beta phosphates, respectively) via ^31^ P NMR chemical shift data (Jancan and Macnaughtan, 2012); thus, above these pKa values, one might expect a larger population of the phosphate-deprotonated UDP-Glc species. Partial charges for UDP and UDP-Glc (UDP moiety only) were obtained from project F-90 of the RESP ESP Charge Database (R.E.DD.B.) (Dupradeau et al., 2008); fragments 1 (“POP”, alpha and beta phosphates for UDP-Glc), 2 (“P1”, alpha phosphate for UDP), 3 (“P1M”, beta phosphate for UDP), and 47 (“U5”, uridine) of this project were used. Partial charges for the glucosyl group of UDP-Glc were obtained from a terminal beta-D-glucose unit of GLYCAM06 (entry “0GB” in its library). However, for the carbon at the reducing end of this glucosyl group linked to the beta phosphate, a charge adjustment of +0.0102 was applied to this glucosyl “C1” atom to provide an integer net charge of -2.0 for the UDP-Glc molecule; within the modular or fragment-based framework of GLYCAM06 and related force fields, such charge adjustments at linking atoms have been used for carbohydrate derivatives, e.g., O-acetyl, O-methyl, O-sulfate, and N-sulfate modifications, and are applicable to other derivatives.

#### MD simulations

The MD simulation protocol employed is based on those used previously to simulate protein and lipid systems (Singh et al., 2020) (Lee et al., 2016; Nixon *et al*., 2016; Sethaphong et al., 2013). Briefly, this protocol involved up to 10,000 steps of energy minimization, gradual NVT heating to 300 K over 100 ps, NVT equilibration at 300 K for 200 ps, NPT equilibration at 300 K and 1 atm for 800 ps, and 1,000 ns of production MD at 300 K and 1 atm. A cutoff of 1.0 nm was used for all stages. The timestep was initially 1.0 fs but was increased to 2.0 fs during the NPT equilibration and NPT production stages. Complete details of the employed protocol, such as the thermostat, barostat, and treatment of long-range interactions are available in a previous report with references cited therein (Singh *et al*., 2020). Above, “NVT” and “NPT” refer to thermodynamic ensembles described by a constant number of particles-fixed volume-regulated temperature and constant number of particles-regulated pressure and temperature, respectively. As noted, AMBER 2021 was used for these present simulations; specifically, AMBER’s pmemd.MPI (CPUs only) and pmemd.cuda (GPU accelerated) were used for the energy minimization and subsequent simulation stages, respectively (Le Grand et al., 2013; Salomon-Ferrer et al., 2013). These simulations were carried out with GPU-equipped servers supplied by Exxact Corporation.

#### MD analysis

Contacts between residues of the CesA8 gating loop (847 to 869) and the ligand groups (uridine, alpha phosphate, beta phosphate, Mg^2+^, and glucosyl group, as applicable) were analyzed using PyContact 1.0.4 (Scheurer *et al*., 2018) with MDAnalysis 0.20.1 (Gowers *et al*., 2016; Michaud-Agrawal *et al*., 2011). Contact criteria included considering heavy atoms only, a contact distance cutoff of 0.5 nm, a hydrogen bond distance cutoff of 0.25 nm, and a hydrogen bond angle cutoff of 120 degrees (Scheurer *et al*., 2018). Contact metrics were calculated and accumulated over atom-atom contacts to obtain residue-residue contact data, where “residue” here can refer to an amino acid residue or a ligand group as listed above. Contact metrics of interest included the mean contact score (based on a distance-weighted sigmoidal function), mean contact lifetime (ns), total contact time (ns and %), and hydrogen bond time (%). The contact analysis was conducted using 1,000 evenly sampled frames from each simulation coordinate trajectory representing 1,000 ns of time.

#### Quantification and Statistical Analysis

Average and standard deviation values were determined using AVERAGE and STDEV functions in Excel and data plotting were performed on GraphPad Prism 6.0.

## Supplemental Information Titles and Legends

**Figure S1. Cryo-EM data processing workflow, related to Figure 1. (A)** UDP bound CesA8. **(B)** *UDP-Glc bound CesA8*.

**Figure S2. Examples of cryo-EM map qualities, related to Figure 1. (A)** UDP-Glc bound CesA8 and **(B)** UDP bound CesA8.

**Figure S3. Comparison of substrate binding poses, related to Figure 1. (A)** Overlay of UDP and UDP-Glc substrate poses. **(B-D) Substrate binding to** *Rhodobacter sphaeroides* BcsA (B, PDB: 5EIY), chitin synthase (C, PDB: 7STM), and hyaluronan synthase (D, PDB: 7SP8).

**Figure S4. Time course of cellulose biosynthesis, related to Figure 2. (A)** Cellulose synthesis reaction for wild-type (WT) BcsA-B IMVs was performed at 37ºC for different time periods starting from 0 to 180 min. At each time interval, 20 µl of reaction mixture was withdrawn, and 2% SDS was added to terminate the synthesis reaction. The products were quantified by scintillation counting. **(B)** Time course of product accumulation for wild-type (WT) poplar CesA8. CesA8 synthesis reactions were incubated at 30ºC and at each indicated time interval, a sample was withdrawn and spotted onto Whatman-2MM blotting paper for quantification. DPM: Disintegrations per minute.

**Figure S5. CesA8’s gating loop interacts with the nucleotide at the active site, related to Figure 3. (A)** Shown is the cryo-EM map of the UDP-Glc-bound CesA8 complex at a low contour level. The green ribbon indicates the gating loop position in UDP-bound *Rhodobacter* BcsA (PDB: 4P00). **(B and C)** Interactions of CesA8’s gating loop with UDP or UDP-Glc and Mg^2+^. A chord diagram based on mean contact scores between gating loop residues of CesA8, groups of UDP/UDP-Glc (URD = uridine, PHA = alpha phosphate, PHB = beta phosphate, and GLC = glucosyl group), and the magnesium ion (MG). The mean contact scores are averaged over time (1,000 ns) using 1,000 evenly sampled frames from each simulation coordinate trajectory. The widths of the nodes (arcs) and links (arrows) are weighted by the mean contact scores, and the links are colored based on their UDP/UDP-Glc/Mg^2+^ destination arcs. (Generated with Origin (Origin)).

**Table S1. EM and model stats, related to Figure 1**.

**Table S2. Contact pairs between CesA protein residues and ligand groups (UDP and Mg**^**2+**^ **), and their contact metrics, related to Figure 3 and S5**. The ligand groups include URD (uridine), PHA (alpha phosphate), PHB (beta phosphate), and MG (Mg^2+^). The contacts are sorted by CesA residue ID number and then by mean score. The numerical columns, mean score to hydrogen bond time, are colored with a green-yellow-red scale from highest to lowest.

**Table S3. Contact pairs between CesA protein residues and ligand groups (UDP-Glc and Mg**^**2+**^ **), and their contact metrics, related to Figure 3 and S5**. The ligand groups include URD (uridine), PHA (alpha phosphate), PHB (beta phosphate), GLC (glucosyl group), and MG (Mg^2+^). The contacts are sorted by CesA residue ID number and then by mean score. The numerical columns, mean score to hydrogen bond time, are colored with a green-yellow-red scale from highest to lowest.

**Table S4. Additional force field parameters employed for UDP-Glc assigned by analogy to GLYCAM06, and modified hydroxyl hydrogen “HO” parameters applied broadly based on that described previously for modified nucleic acids, related to Figure 3 and S5**. The bond parameters include the bond force constant (kcal/mol/Å^2^) and equilibrium bond length (Å). The angle parameters include the angle force constant (kcal/mol/rad^2^) and equilibrium angle (degrees). The dihedral parameters include the energy barrier division factor, half of the energy barrier height (kcal/mol), phase angle (degrees), and the dihedral multiplicity; a negative dihedral multiplicity only indicates that there are additional subsequent terms. The Lennard-Jones (LJ) parameters include half of the interatomic separation distance at the LJ energy minimum, Rmin (Å), and the energy-well depth at the energy minimum, ε (kcal/mol).

