## Supplementary Information for "Insights into substrate coordination and glycosyl transfer of poplar cellulose synthase-8"

### Supplemental Information

Supplemental Figure 1 - 5

Supplemental Table 1 – 5

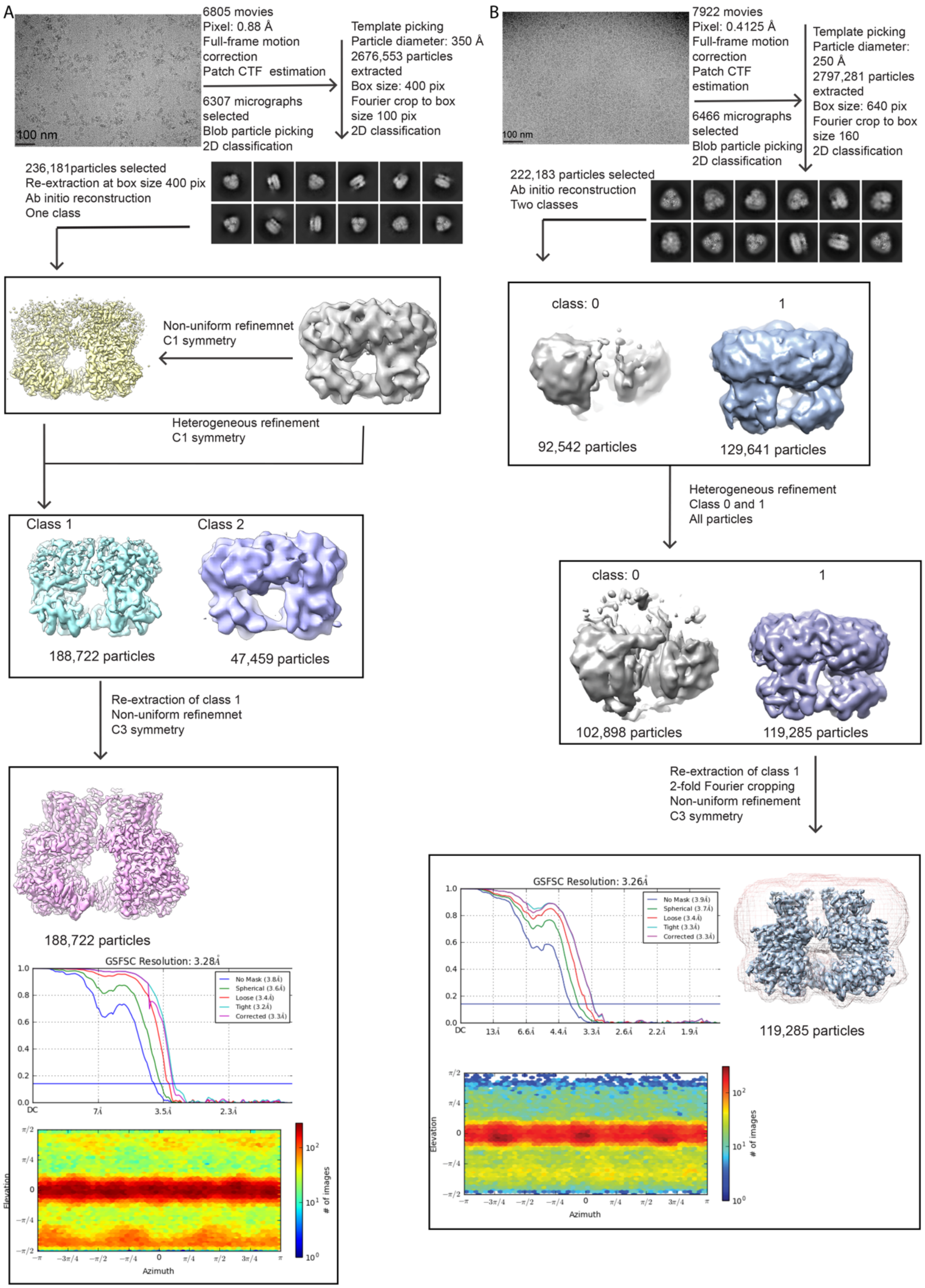

**Figure S1. Cryo-EM data processing workflow, related to Figure 1.** (**A**) UDP bound CesA8. (**B**) UDP-Glc bound CesA8.

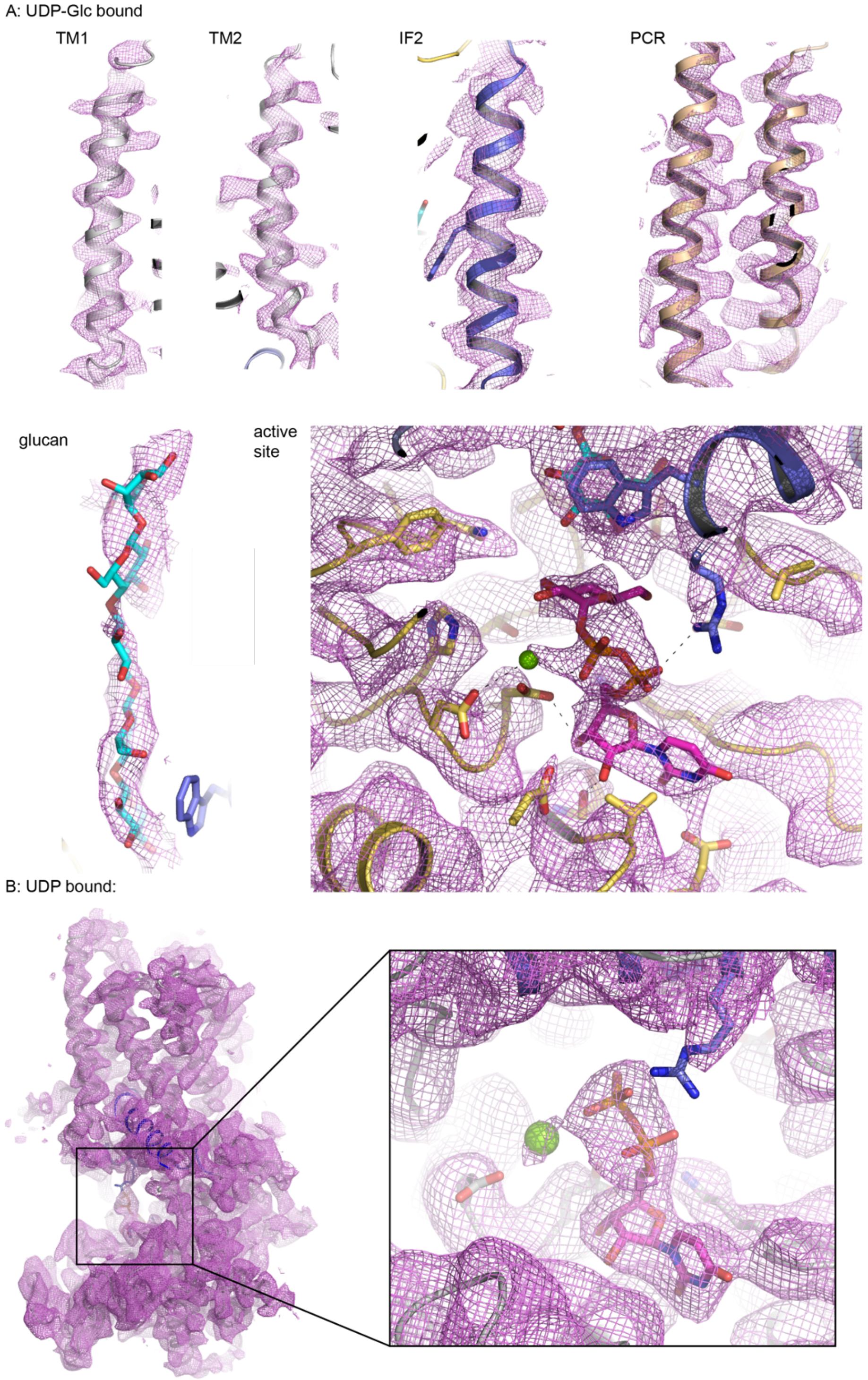

**Figure S2. Examples of cryo-EM map qualities, related to Figure 1.** (**A**) UDP-Glc bound CesA8 and (**B**) UDP bound CesA8.

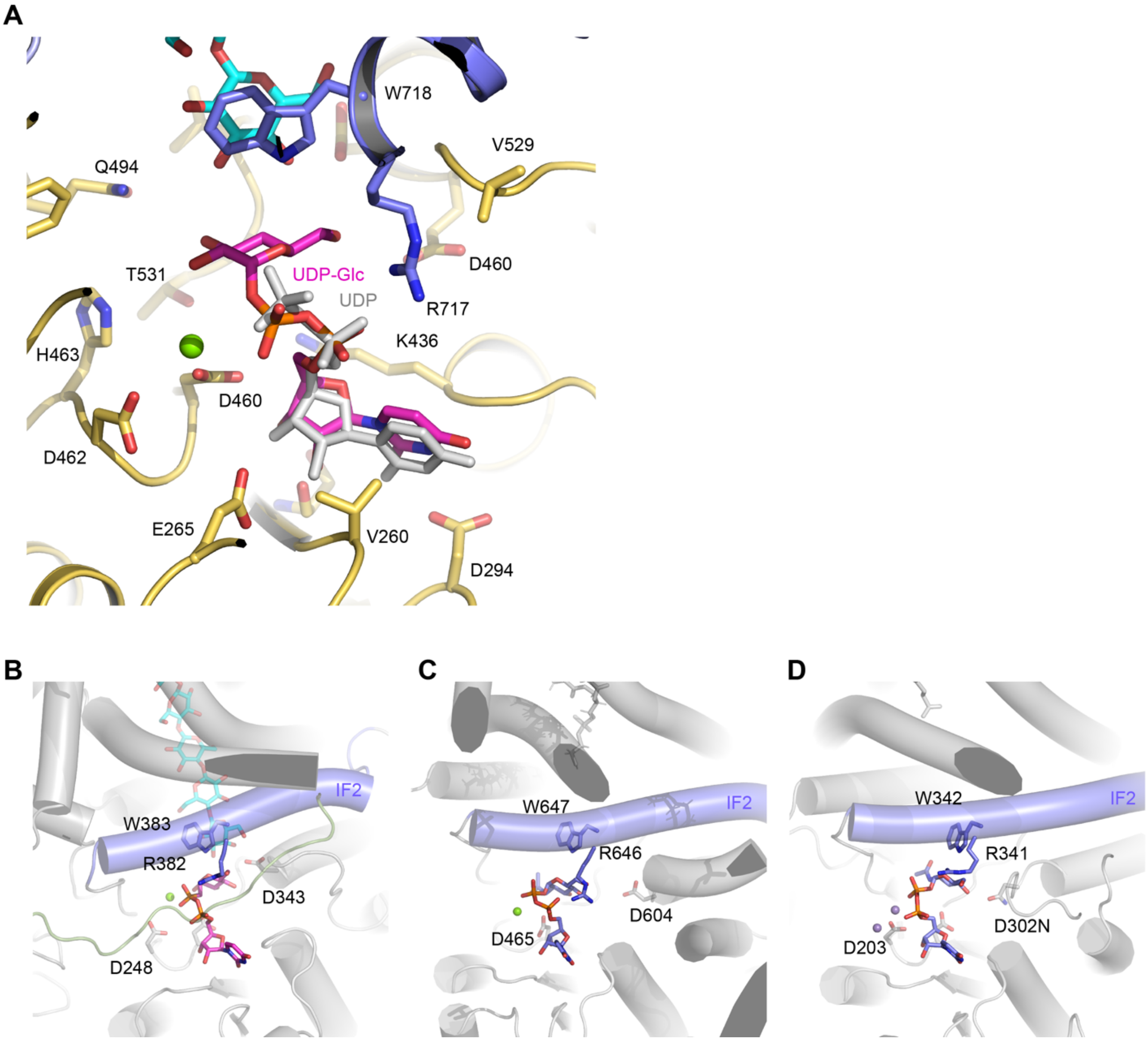

**Figure S3. Comparison of substrate binding poses**, **related to Figure 1. (A)** Overlay of UDP and UDP-Glc substrate poses bound to poplar CesA8. **(B-D) Substrate binding to** *Rhodobacter sphaeroides* BcsA (B, PDB: 5EIY), chitin synthase (C, PDB: 7STM), and hyaluronan synthase (D, PDB: 7SP8).

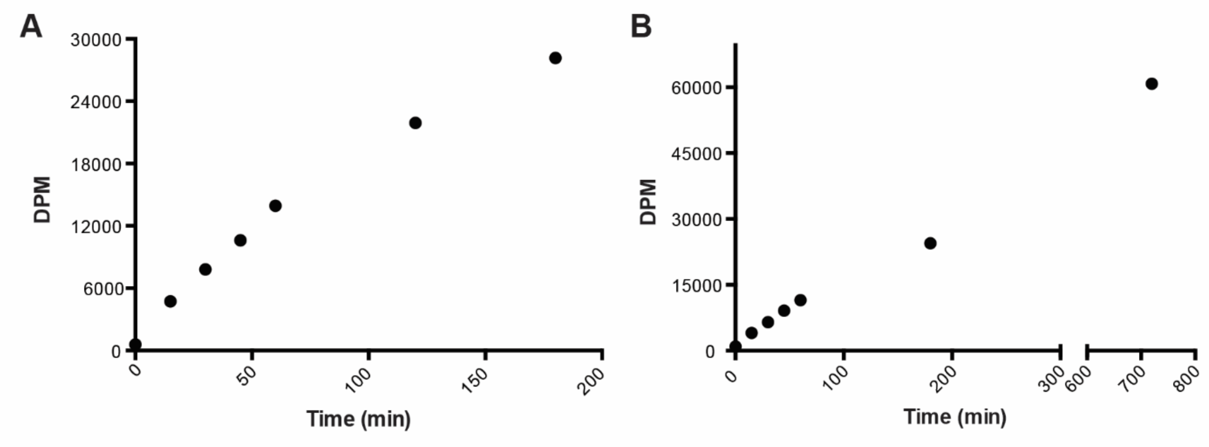

**Figure S4. Time course of cellulose biosynthesis, related to Figure 2. (A)** Cellulose synthesis reaction for wild-type (WT) BcsA-B IMVs was performed at 37ºC for different time periods starting from 0 to 180 min. At each time interval, 20 µl of reaction mixture was withdrawn, and 2% SDS was added to terminate the synthesis reaction. The products were quantified by scintillation counting. **(B)** Time course of product accumulation for wild-type (WT) poplar CesA8. CesA8 synthesis reactions were incubated at 30ºC and at each indicated time interval, a sample was withdrawn and spotted onto Whatman-2MM blotting paper for quantification. DPM: Disintegrations per minute.

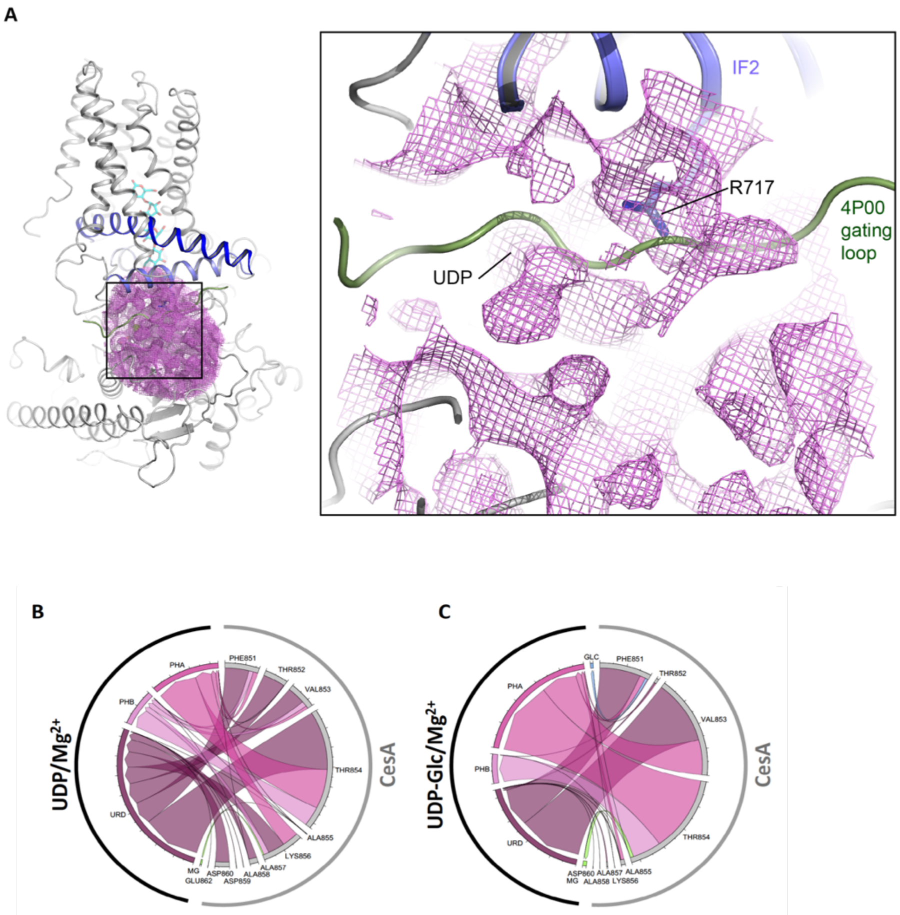

**Figure S5. CesA8's gating loop interacts with the nucleotide at the active site, related to Figure 3.** (**A**) Shown is the cryo-EM map of the UDP-Glc-bound CesA8 complex at a low contour level. The green ribbon indicates the gating loop position in UDP-bound *Rhodobacter* BcsA (PDB: 4P00). (**B** and **C**) Interactions of CesA8's gating loop with UDP or UDP-Glc and $Mg^{2+}$. A chord diagram based on mean contact scores between gating loop residues of CesA8, groups of UDP/UDP-Glc (URD = uridine, PHA = alpha phosphate, PHB = beta phosphate, and GLC = glucosyl group), and the magnesium ion (MG). The mean contact scores are averaged over time (1,000 ns) using 1,000 evenly sampled frames from each simulation coordinate trajectory. The widths of the nodes (arcs) and links (arrows) are weighted by the mean contact scores, and the links are colored based on their UDP/UDP-Glc/$Mg^{2+}$ destination arcs. (Generated with Origin (Origin)).

### Table S1. EM and model stats, related to Figure 1.

| Cryo-electron microscopy data collection and processing | | |
|---|---|---|
| | **UDP-Glc bound** | **UDP bound** |
| Microscope | FEI Titan Krios G3i | FEI Titan Krios G3i |
| Voltage (keV) | 300 | 300 |
| Camera | Gatan K3 | Gatan K3 |
| Energy Filter | BioQuantum | BioQuantum |
| Pixel size (Å) | 0.412 | 0.88 |
| Defocus range (µm) | -2.3 – -0.8 | -2.3 – -0.8 |
| Magnification (nominal) | 105,000 | 81,000 |
| Electron exposure ($e^-$/pix/s) | 15 | 15 |
| Exposure rate ($e^-/Å^2$) | 51 | 50 |
| Frames per movie | 50 | 40 |
| Energy filter slit width (eV) | 20 | 10 |
| Automation software | EPU | EPU |
| Micrographs used | 6466 | 6307 |
| Extracted particles | 2,797,281 | 2,676,553 |
| Particles in final 3D refinement | 119,285 | 188,722 |
| Resolution No mask (Å) | 3.9 | 3.8 |
| Resolution Spherical | 3.7 | 3.6 |
| Resolution Loose | 3.4 | 3.4 |
| Resolution Tight | 3.3 | 3.3 |
| Resolution Corrected | 3.3 | 3.3 |
| Sharpening B-factor (Å2) | -130 | -129 |
| EMDB ID | | |
| **Coordinate Refinement and Validation** | | |
| Refinement program | Phenix (Real-space refinement) | Phenix (Real-space refinement) |
| Number of protein atoms (non-H) | 34467 | 34562 |
| Number of ligands | CE5: UDP-Glc:MG3:3:3 | CE5:UDP:Mg 3:3:3 |
| RMSD bond (Å) | 0.002 | 0.002 |
| RMSD angle (°) | 0.595 | 0.562 |
| Ramachandran favored (%) | 95.14 | 94.62 |
| Ramachandran allowed (%) | 4.68 | 5.19 |
| Ramachandran outlier (%) | 0.19 | 0.19 |
| All-atom clash score | 8.67 | 5.06 |
| MolProbity Score | 1.8 | 1.64 |
| B-factors (min/max/mean) | | |
| Protein | 34/153/82 | 8/211/92 |
| Overall correlation coefficient | | |
| CC (mask) | 0.75 | 0.83 |
| CC (box) | 0.59 | 0.68 |
| CC (peaks) | 0.49 | 0.6 |
| CC (volume) | 0.73 | 0.81 |
| Mean CC for ligands | 0.6 | 0.67 |
| PDB ID | 8G2J | 8G27 |

**Table S2. Contact pairs between CesA protein residues and ligand groups (UDP and $Mg^{2+}$), and their contact metrics, related to Figure 3 and S5.** The ligand groups include URD (uridine), PHA (alpha phosphate), PHB (beta phosphate), and MG ($Mg^{2+}$). The contacts are sorted by CesA residue ID number and then by mean score. The numerical columns, mean score to hydrogen bond time, are colored with a green-yellow-red scale from highest to lowest.

| Residue | Ligand group | Mean score | Mean lifetime (ns) | Total time (ns) | Total time (%) | Hydrogen bond time (%) |
|---|---|---|---|---|---|---|
| PHE851 | URD | 0.84 | 73.7 | 221.0 | 22.1 | 0.0 |
| PHE851 | PHA | 0.00 | 1.6 | 61.0 | 6.1 | 0.0 |
| THR852 | PHA | 0.17 | 24.5 | 955.0 | 95.5 | 0.0 |
| THR852 | URD | 0.05 | 4.5 | 303.0 | 30.3 | 0.0 |
| THR852 | PHB | 0.00 | 0.5 | 1.0 | 0.1 | 0.0 |
| VAL853 | PHA | 4.43 | 1000.0 | 1000.0 | 100.0 | 0.0 |
| VAL853 | URD | 1.99 | 248.5 | 994.0 | 99.4 | 0.0 |
| VAL853 | PHB | 0.00 | 1.4 | 7.0 | 0.7 | 0.0 |
| THR854 | PHA | 7.83 | 1000.0 | 1000.0 | 100.0 | 99.9 |
| THR854 | PHB | 3.39 | 1000.0 | 1000.0 | 100.0 | 24.8 |
| THR854 | MG | 0.06 | 15.6 | 172.0 | 17.2 | 0.0 |
| THR854 | URD | 0.00 | 1.2 | 23.0 | 2.3 | 0.0 |
| ALA855 | PHA | 2.37 | 75.8 | 986.0 | 98.6 | 34.5 |
| ALA855 | URD | 0.19 | 6.8 | 506.0 | 50.6 | 0.0 |
| ALA855 | PHB | 0.14 | 2.3 | 337.0 | 33.7 | 0.0 |
| ALA855 | MG | 0.00 | 1.9 | 71.0 | 7.1 | 0.0 |
| LYS856 | PHB | 0.10 | 3.9 | 27.0 | 2.7 | 1.4 |
| LYS856 | MG | 0.01 | 2.5 | 25.0 | 2.5 | 0.0 |
| ALA857 | URD | 0.01 | 1.2 | 31.0 | 3.1 | 0.0 |
| ALA858 | URD | 0.65 | 12.1 | 555.0 | 55.5 | 0.0 |
| ASP859 | URD | 0.07 | 1.9 | 106.0 | 10.6 | 0.8 |
| ASP860 | URD | 0.01 | 1.3 | 16.0 | 1.6 | 0.0 |
| THR861 | URD | 0.00 | 0.5 | 1.0 | 0.1 | 0.0 |

**Table S3. Contact pairs between CesA protein residues and ligand groups (UDP-Glc and $Mg^{2+}$), and their contact metrics, related to Figure 3 and S5**. The ligand groups include URD (uridine), PHA (alpha phosphate), PHB (beta phosphate), GLC (glucosyl group), and MG ($Mg^{2+}$). The contacts are sorted by CesA residue ID number and then by mean score. The numerical columns, mean score to hydrogen bond time, are colored with a green-yellow-red scale from highest to lowest.

| Residue | Ligand group | Mean score | Mean lifetime (ns) | Total time (ns) | Total time (%) | Hydrogen bond time (%) |
|---|---|---|---|---|---|---|
| ASN850 | URD | 0.00 | 0.5 | 1.0 | 0.1 | 0.0 |
| PHE851 | URD | 2.80 | 64.7 | 841.0 | 84.1 | 0.0 |
| PHE851 | GLC | 1.15 | 12.1 | 653.0 | 65.3 | 0.0 |
| PHE851 | PHA | 0.08 | 3.7 | 467.0 | 46.7 | 0.0 |
| PHE851 | PHB | 0.00 | 1.8 | 97.0 | 9.7 | 0.0 |
| THR852 | PHA | 0.10 | 13.0 | 844.0 | 84.4 | 0.0 |
| THR852 | URD | 0.01 | 1.3 | 52.0 | 5.2 | 0.0 |
| THR852 | GLC | 0.00 | 1.0 | 5.0 | 0.5 | 0.0 |
| THR852 | PHB | 0.00 | 0.5 | 1.0 | 0.1 | 0.0 |
| VAL853 | URD | 5.78 | 1000.0 | 1000.0 | 100.0 | 0.0 |
| VAL853 | PHA | 4.14 | 1000.0 | 1000.0 | 100.0 | 0.0 |
| VAL853 | PHB | 0.00 | 0.8 | 4.0 | 0.4 | 0.0 |
| VAL853 | MG | 0.00 | 0.7 | 2.0 | 0.2 | 0.0 |
| THR854 | PHA | 7.20 | 1000.0 | 1000.0 | 100.0 | 99.9 |
| THR854 | PHB | 3.46 | 1000.0 | 1000.0 | 100.0 | 54.6 |
| THR854 | MG | 0.19 | 9.5 | 815.0 | 81.5 | 0.0 |
| THR854 | URD | 0.01 | 2.2 | 161.0 | 16.1 | 0.0 |
| ALA855 | PHA | 2.26 | 22.2 | 845.0 | 84.5 | 50.8 |
| ALA855 | URD | 0.55 | 11.9 | 451.0 | 45.1 | 0.0 |
| ALA855 | MG | 0.06 | 11.6 | 671.0 | 67.1 | 0.0 |
| ALA855 | PHB | 0.00 | 1.7 | 73.0 | 7.3 | 0.0 |
| LYS856 | URD | 0.02 | 1.7 | 77.0 | 7.7 | 0.2 |
| LYS856 | MG | 0.01 | 2.4 | 352.0 | 35.2 | 0.0 |
| LYS856 | PHA | 0.01 | 2.1 | 199.0 | 19.9 | 0.0 |
| ALA858 | URD | 0.01 | 1.7 | 54.0 | 5.4 | 0.0 |
| ASP860 | URD | 0.24 | 6.3 | 339.0 | 33.9 | 0.0 |

**Table S4. Additional force field parameters employed for UDP-Glc assigned by analogy to GLYCAM06, and modified hydroxyl hydrogen "HO" parameters applied broadly based on that described previously for modified nucleic acids, related to Figure 3 and S5**. The bond parameters include the bond force constant (kcal/mol/Å$^2$) and equilibrium bond length (Å). The angle parameters include the angle force constant (kcal/mol/rad$^2$) and equilibrium angle (degrees). The dihedral parameters include the energy barrier division factor, half of the energy barrier height (kcal/mol), phase angle (degrees), and the dihedral multiplicity; a negative dihedral multiplicity only indicates that there are additional subsequent terms. The Lennard-Jones (LJ) parameters include half of the interatomic separation distance at the LJ energy minimum, Rmin (Å), and the energy-well depth at the energy minimum, ε (kcal/mol).

| Name | Atom types & parameters | Source |
|---|---|---|
| Bond 1 | Cg-OS 285.00 1.46 | GLYCAM_06j.dat |
| Angle 1 | H2-Cg-OS 60.00 110.00 | GLYCAM_06j.dat |
| Angle 2 | Os-Cg-OS 100.00 112.00 | GLYCAM_06j.dat |
| Angle 3 | OS-Cg-Cg 70.00 108.50 | GLYCAM_06j.dat |
| Angle 4 | P -OS-Cg 50.0 118.88 | GLYCAM_06j.dat |
| Dihedral 1 | OS-Cg-Os-Cg 1 0.96 0.0 -3.<br>1 1.38 0.0 -2.<br>1 1.08 0.0 1. | GLYCAM_06j.dat |
| Dihedral 2 | OS-Cg-Cg-Cg 1 -0.27 0.0 1. | GLYCAM_06j.dat |
| Dihedral 3 | H1-Cg-Cg-OS 1 0.05 0.0 3. | GLYCAM_06j.dat |
| Dihedral 4 | Oh-Cg-Cg-OS 1 -1.10 0.0 -1.<br>1 0.25 0.0 2. | GLYCAM_06j.dat |
| Dihedral 5 | H2-Cg-OS-P 1 0.17 0.0 3. | GLYCAM_06j.dat |
| Dihedral 6 | Os-Cg-OS-P 1 -1.20 0.0 1. | GLYCAM_06j.dat |
| Dihedral 7 | Cg-Cg-OS-P 1 -1.20 0.0 1. | GLYCAM_06j.dat |
| LJ 1 | HO 0.6000 0.0157 | all_modrna08.frcmod |

**Table S5. Oligonucleotides used for generating BcsA and CesA8 mutants, related to Star Methods.** Mutated sequences are indicated with lowercase letters.

| Name | Oligonucleotides |
|---|---|
| BcsA-R382F-Fw | CCTTCATCCAGCAGCGCGGCttcTGGGCCACCGGCATGATGCAG |
| BcsA-R382F-Rv | CTGCATCATGCCGGTGGCCCAgaaGCCGCGCTGCTGGATGAAGG |
| BcsA-R382A-Fw | TGGGCGACGGGTATGATGCAGATGCTGCTGCTGAAG |
| BcsA-R382A-Rv | CGTCGCCCAcgcGCCGCGCTGCTGGATGAAG |
| BcsA-F503I-Fw | GCCGTTACTGCCAAGGACGAGACGCTGAGCGAG |
| BcsA-F503I-Rv | AGTAACGGCaatGCGGGCACTGCGCGGCCGCAG |
| BcsA-F503A-Fw | TGCGGCCGCGCAGTGCCCGCgccGCGGTGACCGCGAAGGACGAGAC |
| BcsA-F503A-Rv | GTCTCGTCCTTCGCGGTCACCGCggcGCGGGCACTGCGCGGCCGCA |
| BcsA-V505L-Fw | CCGCGCAGTGCCCGCTTCGCGctgACCGCGAAGGACGAGACGCTG |
| BcsA-V505L-Rv | CAGCGTCTCGTCCTTCGCGGTcagCGCGAAGCGGGCACTGCGCGG |
| BcsA-V505A-Fw | GCGCAGTGCCCGCTTCGCGgcgACCGCGAAGGACGAGACGCTG |
| BcsA-V505A-Rv | CAGCGTCTCGTCCTTCGCGGTcgcCGCGAAGCGGGCACTGCGC |
| BcsA-T506S-Fw | CGCAGTGCCCGCTTCGCGGTGtccGCGAAGGACGAGACGCTGAG |
| BcsA-T506S-Rv | CTCAGCGTCTCGTCCTTCGCggaCACCGCGAAGCGGGCACTGCG |
| BcsA-T506A-Fw | CGCAGTGCCCGCTTCGCGGTGgcgGCGAAGGACGAGACGCTGAGCG |
| BcsA-T506A-Rv | CGCTCAGCGTCTCGTCCTTCGCcgcCACCGCGAAGCGGGCACTGCG |
| BcsA-K508R-Fw | GCTTCGCGGTGACCGCGcgcGACGAGACGCTGAGCGAGAAC |
| BcsA-K508R-Rv | GTTCTCGCTCAGCGTCTCGTCgcgCGCGGTCACCGCGAAGC |
| BcsA-K508A-Fw | GATGAAACCTTAAGCGAGAACTACATTTCG |
| BcsA-K508A-Rv | GGTTTCATCggcCGCGGTCACCGCGAAGCGGGC |
| CesA8-R717A-Fw | CACCAGGTTCTCgcaTGGGCTCTTGGA |
| CesA8-R717A-Rv | TCCAAGAGCCCAtgcGAGAACCTGGTG |
| CesA8-F851I-Fw | ATTGATACGAACattACTGTCACAGCA |
| CesA8-F851I-Rv | TGCTGTGACAGTaatGTTCGTATCAAT |
| CesA8-V853L-Fw | ACGAACTTTACTctcACAGCAAAAGCA |
| CesA8-V853L-Rv | TGCTTTTGCTGTgagAGTAAAGTTCGT |
| CesA8-T854S-Fw | AACTTTACTGTCtcaGCAAAAGCAGCC |
| CesA8-T854S-Rv | GGCTGCTTTTGCtgaGACAGTAAAGTT |
| CesA8-T854A-Fw | AACTTTACTGTCgcaGCAAAAGCAGCC |
| CesA8-T854A-Rv | GGCTGCTTTTGCtgcGACAGTAAAGTT |
| CesA8-K856R-Fw | ACTGTCACAGCAagaGCAGCCGATGAT |
| CesA8-K856R-Rv | ATCATCGGCTGCtctTGCTGTGACAGT |
